# Characterization of Blood Group Variants in an Omani Population by Comparison of Whole Genome Sequencing and Serology

**DOI:** 10.1101/2024.06.17.599396

**Authors:** Paige E. Haffener, Arwa Z. Al-Riyami, Shoaib Al-Zadjali, Mohammed Al-Rawahi, Saif Al Hosni, Ali Al Marhoobi, Ammar Al Sheriyani, Ellen M. Leffler

**Affiliations:** Department of Human Genetics, The University of Utah School of Medicine, Salt Lake City, UT, USA; Department of Hematology, Sultan Qaboos University Hospital, University Medical City, Muscat, Oman; Sultan Qaboos Comprehensive Cancer Center, University Medical City, Muscat, Oman; Department of Hematology, College of Medicine and Health Sciences, Sultan Qaboos University, Muscat, Oman; Royal Oman Police Hospital, Muscat, Oman

## Abstract

Although blood group variation was first described over a century ago, our understanding of the genetic variation affecting antigenic expression on the red blood cell surface in many populations is lacking. This deficit limits the ability to accurately type patients, especially as serological testing is not available for all described blood groups, and targeted genotyping panels may lack rare or population-specific variants. Here, we perform serological assays across 24 antigens and whole genome sequencing on 100 Omanis, a population underrepresented in genomic databases. We inferred blood group phenotypes using the most commonly typed genetic variants. The comparison of serological to inferred phenotypes resulted in an average concordance of 96.9%. Among the 22 discordances, we identify seven known variants in four blood groups that, to our knowledge, have not been previously reported in Omanis. Incorporating these variants for phenotype inference, concordance increases to 98.8%. Additionally, we describe five candidate variants in the Lewis, Lutheran, MNS, and P1 blood groups that may affect antigenic expression, although further functional confirmation is required. Notably, we identify several blood group alleles most common in African populations, likely introduced to Oman by gene flow over the last thousand years. These findings highlight the need to evaluate individual populations and their population history when considering variants to include in genotype panels for blood group typing. This research will inform future work in blood banks and transfusion services.

**Key Points:**

- Utilizing whole genome sequencing to infer blood types in Omanis demonstrates high sensitivity for most blood groups
- Population history influences blood group variation, necessitating population-specific genotype panels

## Introduction

In the 124 years since the discovery of the ABO blood group system, 45 different blood groups have been described in humans along with 50 associated blood group genes^1^. Despite extensive knowledge of these various blood group systems, most have been described via case studies, and recent population genomic analyses suggest there is much left to be discovered. For instance, a study of genomic variation in African, European, South Asian, East Asian, and American populations from the 1000 Genomes project^2^ identified 1,241 nonsynonymous (NS) variants within 43 blood groups genes, and reported that 1,000 of the NS variants (81%) were not known blood group polymorphisms, yet 357 were extracellular and thus potentially antigenic^3^. Another study of the same dataset identified only 120 of 604 known blood group variants^4^, with 36 of these found in at least one continental region where they had not previously been described^4^. This suggests that many known variants affecting blood group variation are rare, and that we still lack a complete understanding of the distribution of blood group alleles in many populations^4^.

For example, a recent study from Oman demonstrated that rare, undescribed variants likely affect antigenic expression in multiple blood groups^5^. In this study, targeted genotyping and serological assays were compared for 19 different antigens belonging to six blood group systems. Although overall concordance was high (>95%), Fy^b^ was an exception (concordance 87%), and only 3 antigens had 100% concordance^5^. Discordances were likely due to the effect of genetic variants that were absent from the genotyping assay. While the prevalence of common blood group antigens or blood group alleles have been documented across much of the Arabian Peninsula^6,7,8-11,12^, no comparison of sequencing data and serology has been conducted to identify additional variants affecting antigenic expression in this region.

Targeted genotyping is increasingly being investigated as an alternative or complement to serology for blood group phenotyping^13^. Benefits of genotyping include improving red cell matching for multi-transfused patients, those at an increased risk of alloimmunization, such as patients with sickle cell disease, and those with autoantibodies. The addition of a genotyping strategy is of interest in Oman where hemoglobinopathies and risk of alloimmunization are common in the population^14-16^. However, a more complete picture of rare and population-specific variants affecting antigenic expression is necessary. Whole genome sequencing provides a comprehensive view of blood group loci, including indels and copy number variation, particularly at more complex loci such as those that determine the RH and MNS blood groups^17,18^.

Here, we compare antigen typing inferred by whole genome sequencing to antigen expression determined by serology to identify variants contributing to blood group variation in the Omani population. We identified seven variants that have previously not been described in Omanis. Additionally, we identified five candidate variants that may be affecting antigen expression in the Lewis, Lutheran, MNS and P1 blood groups by altering erythrocyte-specific transcription factor binding sites, or by altering the coding regions near alleles encoding the blood group antigens. These findings should be considered when selecting red cell genotyping platforms for blood banks and transfusion services in Oman and nearby regions.

## Methods

### Sample Collection

A description of the samples used in this analysis, including DNA extraction and shipping conditions, has been previously published^19^. Briefly, 100 healthy male and female Omani blood donors between the ages of 18 and 60 years attending the Sultan Qaboos University Hospital (SQUH) blood bank were randomly selected and consented for enrollment in the study. The Medical Research Ethics committee at the College of Medicine and Health Sciences, the Sultan Qaboos University approved this study (MREC #2034, 2019).

### Blood Bank Methods

Red blood cell phenotyping was performed within 24 hours of collection at SQUH Blood bank using BioRad© antisera on freshly drawn samples per the manufacturer instructions (BioRad©, Cressier Switzerland) and as previously published^12^. The following blood systems and antigens were tested: ABO (A,B antigens), Rh (C,c,E,e antigens), Kell (K, k, Kp^a^, Kp^b^ antigens), Kidd (Jk^a^, Jk^b^ antigens), Duffy (Fy^a^,Fy^b^ antigens), Lewis (Le^a^,Le^b^ antigens), Lutheran (Lu^a^,Lu^b^ antigens), and MNS (M,N,S,s antigens). A clear red cell button at the bottom of the phenotyping well was defined as a negative reaction for all antigens (grade 0). Rh D reactions of 0 or 1 are further tested for weak D. Reactions positive for weak D are reported as Rh D positive and reactions negative for weak D testing are reported as Rh D negative as per manufacturer instructions. Other reaction patterns were defined as positive and were graded (1-4) for each antigen phenotyped. We included known positive and negative samples as internal controls for each antigen.

### Genome Sequencing, Alignment and Variant Calling

As previously described^19^, short read (150bp paired-end) whole genome sequencing was performed to an average coverage of 16X at the Huntsman Cancer Institute High-Throughput Genomics Shared Resource at the University of Utah. The sequence reads were aligned to GRCh38 with BWA-MEM^20^ and variants were called following the GATK best practices protocol^21,22^. Haplotype phasing was done using Eagle v2^23^ to produce a haplotype variant call file.

### Inferring Blood Group Phenotypes

#### ABO, RHCE, Kell, Kidd, Duffy, Lewis, Lutheran, MNS, and P1 Inference with SNVs

Using the databases available from ISBT^1^ and BloodAntigens.com^13^, we curated a list of variants for inferring blood group phenotypes from SNV genotypes (Table 1).

**Table 1.**
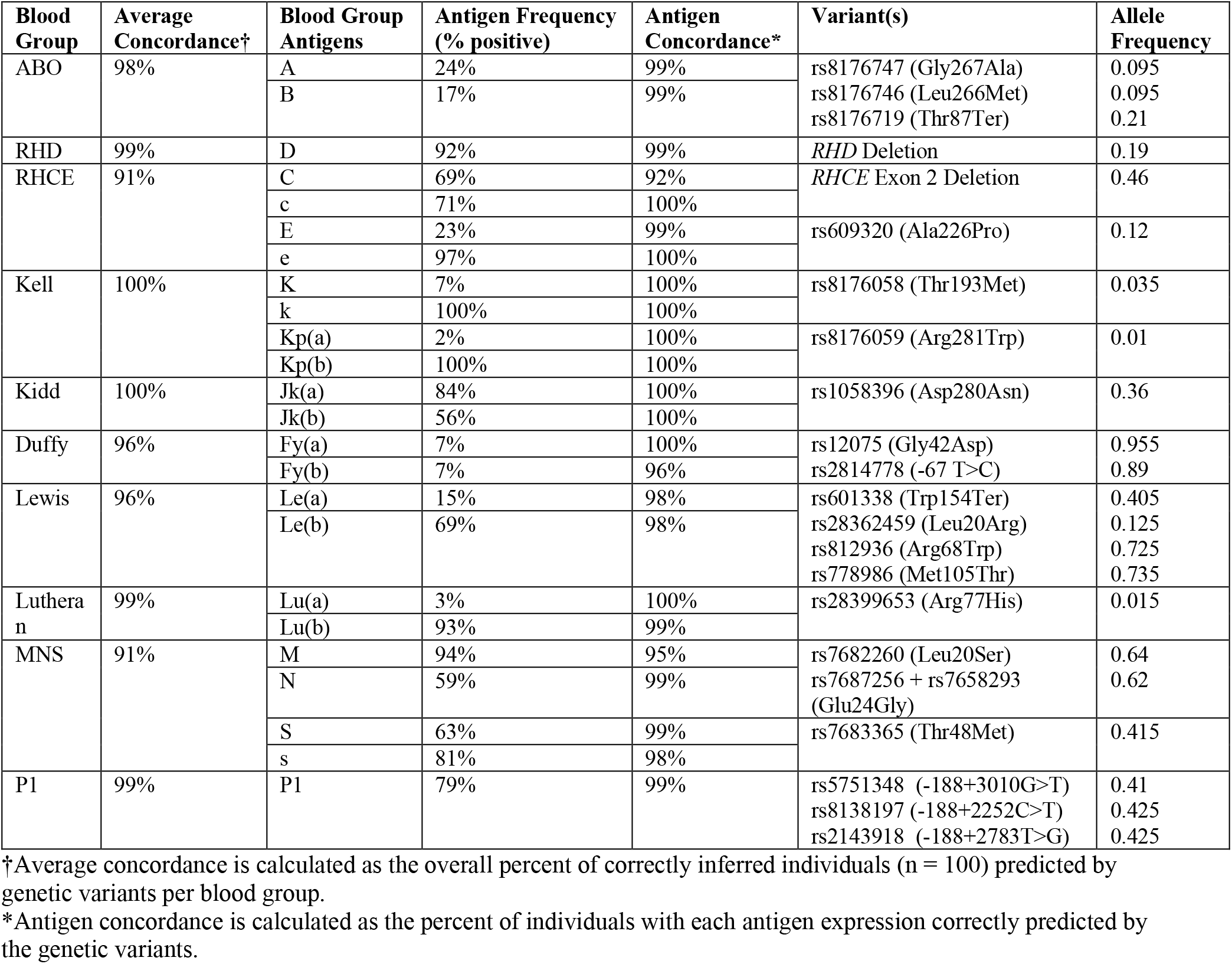
Blood group system and antigen concordances from the comparison of blood typing by serology and inference from the commonly typed genetic variants. Variant information and allele frequency in this dataset are shown in the last two columns.

#### RHD and RHCE Copy Number Inference

RHD phenotypes were inferred using a copy number analysis^13^. Using aligned reads filtered for a MAPQ > 20, coverage across the *RHD* locus (1:25272393-25330445) and *RHCE* locus (1:25360659-25430193) was calculated using SAMtools^24^. Using the equation described by Lane et al. 2018^13^, a ratio of *RHD* to *RHCE* coverage between 0-0.5 was classified as null, 0.6-1.5 as hemizygous, and 1.6-2.5 as homozygous. RHCE C/c antigen phenotypes were also inferred using a copy number analysis suggested by Lane et al. ^13^, comparing coverage of *RHCE* exon 2 to the entire *RHCE* locus. A ratio of coverage greater than or equal to 1.5 was inferred as C-c+, 0.5-1.4 as C+c+, and less than 0.5 as C+c-.

#### MNS Copy Number Inference

To identify copy number variation, we inferred the underlying copy number state from observed coverage at sites with high mappability in 1600bp windows across the *GYPA, GYPB* and *GYPE* loci using a Hidden Markov Model as previously described^19,25^.

#### Concordance calculations

We calculated two concordances in this analysis, considering the phenotype determined by serology as truth. The average concordance per blood group is defined as the overall percent of all correctly inferred phenotypes from genotype data for that blood group. For instance, for the Kidd blood group, this would be calculated as follows:

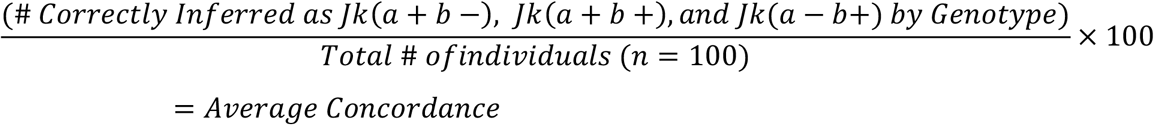

The antigen concordance is calculated per antigen and is defined as the percent of individuals with individual antigen expression correctly predicted by genotype inferences over the total number of individuals. Using the Kidd blood group as an example, antigen concordance for the Jk(a) antigen would be calculated as follows:

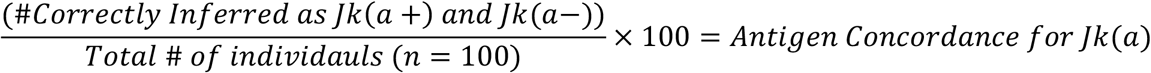

## Results

In 100 Omani blood donors, we compared blood group phenotypes determined by serology (Table S1) and by inference from genetic variants called from whole genome sequencing data for a commonly used serology panel including ABO, Rh, Kell, Kidd, Duffy, Lewis, Lutheran, P1, and MNS (Table 1). Using the most common variants underlying the 24 antigens tested, we evaluated the concordance for each blood group system as well as for each antigen (Table 1). Across all blood group systems, the average concordance was 96.9% (Table 1). Two blood group systems had a phenotype-genotype concordance of 100% (Kell and Kidd). The remaining seven blood groups had a concordance greater than 95% with the exception of the MNS and RHCE C/c blood group systems which had concordances of 91%.

We identified a total of 22 discordant samples (Table S2). Among these, we identified seven known variants in eleven samples affecting antigen expression in the Rh, Duffy, Lewis, and MNS blood groups that were previously undescribed in the Omani population (Table 2). We also identified a putatively novel variant in the Lewis blood group as well as three variants in transcription factor binding sites specific to erythrocyte expression or erythropoiesis that could be altering antigen expression in the Lutheran and P1 blood groups (Table 3). Additionally, we identified a structural variant in the glycophorin gene region that may alter S antigen expression of the MNS blood group (Table 3). There are eight discordances without a candidate novel variant that remain unresolved, all of which are in the Rh and MNS blood groups. We discuss the discordances and the identified variants for each blood group in detail below.

**Table 2.** Summary of known variants resolving discrepancies between inferred phenotypes from whole genome sequence data and phenotypes reported by serology in the Omani samples.

| Blood Group | Phenotype by Serology | Inferred Phenotype | Resolving Variant | Allele Frequency | Number of Discrepancies |
| --- | --- | --- | --- | --- | --- |
| Rh | D- | D+ | RHD*ψ<sup>28</sup> rs748783394 | 0.0376 | 1 |
| Rh | E+e+ | E-e+ | rs141398055 (Arg201Thr) | 0.015 | 1 |
| Duffy | Fy(a-b-) | Fy(a-b+) | Duffy X rs34599082 (Arg89Cys)<sup>19</sup> | 0.015 | 3 |
| Duffy | Fy(a-b-) | Fy(a-b+) | rs773692057 (Ser62fs)<sup>19</sup> | 0.005 | 1 |
| Lewis | Le(a-b-) | Le(a-b+) | rs3894326 (Ile356Lys) | 0.065 | 2 |
| Lewis | Le(a-b-) | Le(a+b-) | rs3894326 (Ile356Lys) | 0.065 | 1 |
| MNS | S-s- | S+s- | GYP.He(P2)<sup>50</sup> rs139511876; DEL1<sup>25</sup> | 0.01; 0.015 | 1 |

**Table 3.** Summary of putatively novel causal variants identified in the discordant Omani samples.

| Blood Group | Phenotype by Serology | Inferred Phenotype | Candidate Variants | Variant Effect | Allele Frequency | Number of Discrepancies |
| --- | --- | --- | --- | --- | --- | --- |
| Lewis | Le(a+b+) | Le(a-b+) | rs373779096 | Amino acid change | 0.005 | 1 |
| Lutheran | In(Lu) | Lu(a-b+) | rs533045163 and rs184739796 | GATA1 binding site | 0.005 and 0.005 | 1 |
| MNS | S+s- | S+s+ | Dantu Structural variant | GYPA – GYPB copy number variation | 0.005 | 1 |
| P1 | P2 | P1 | Chr22:42,721,266 (A4GALT, -289A>C) | STAT1 binding site | 0.01 | 1 |

### ABO Blood Group

The AB and O antigens are encoded by the *ABO* gene on chromosome 9. Using rs8176747 (Gly267Ala) and rs8176746 (Leu265Met) to infer the A and B antigens and rs8176719 (Thr87AspfsTer107) to infer the O antigen resulted in a concordance of 98%. Because Thr87AspfsTer107 is most commonly found on a haplotype that expresses the A antigen^26^, 13 individuals heterozygous for all three SNVs were inferred as blood type B. However, the variants are not in complete linkage disequilibrium (D’=0.8) and one was found to express A by serology. The variants are too far apart for physical phasing, but this sample was imputed as carrying 87AspfsTer107 on the same haplotype as the alleles encoding Ala and Met, consistent with the A blood type. The other discordant sample was called as homozygous for Thr87AspfsTer107 and thereby inferred as O blood type but expressed the B antigen via serology. Further investigation revealed that this individual had one read with the insertion. Sanger sequencing confirmed that this individual was in fact heterozygous for Thr87AspfsTer107, resolving this discordance.

### Rh Blood Group

For the Rh blood group, we typed the D, C, c, E, and e antigens encoded by the adjacent *RHD* and *RHCE* genes. The presence of the *RHD* gene on chromosome 1 results in the expression of D antigen whereas homozygosity for a complete *RHD* gene deletion is the most common cause of the Rh D negative phenotype^27^. Serology and genotype inference were discordant for the D antigen in one individual. This sample was inferred as D+ by sequence data, supported by numerous reads mapping to the *RHD* gene but serologically, the D antigen was not detected. This individual was found to carry the *RHD* pseudogene allele (*RHD*\*ψ) that consists of a 37bp duplication in exon 4, which introduces a premature stop codon resulting in early truncation of the *RHD* gene^28^. The allele frequency (AF) of *RHD*\*ψ in the Omanis is similar to the AF in African/African American population in gnomADv4.0^29^ (Figure 1), with AFs of 0.0376 and 0.0389, respectively. This is higher than the gnomADv4.0 Middle Eastern population (AF=0.0022), indicating heterogeneity across the Middle East, likely due to variation in African ancestry^19^.

**Figure 1.**
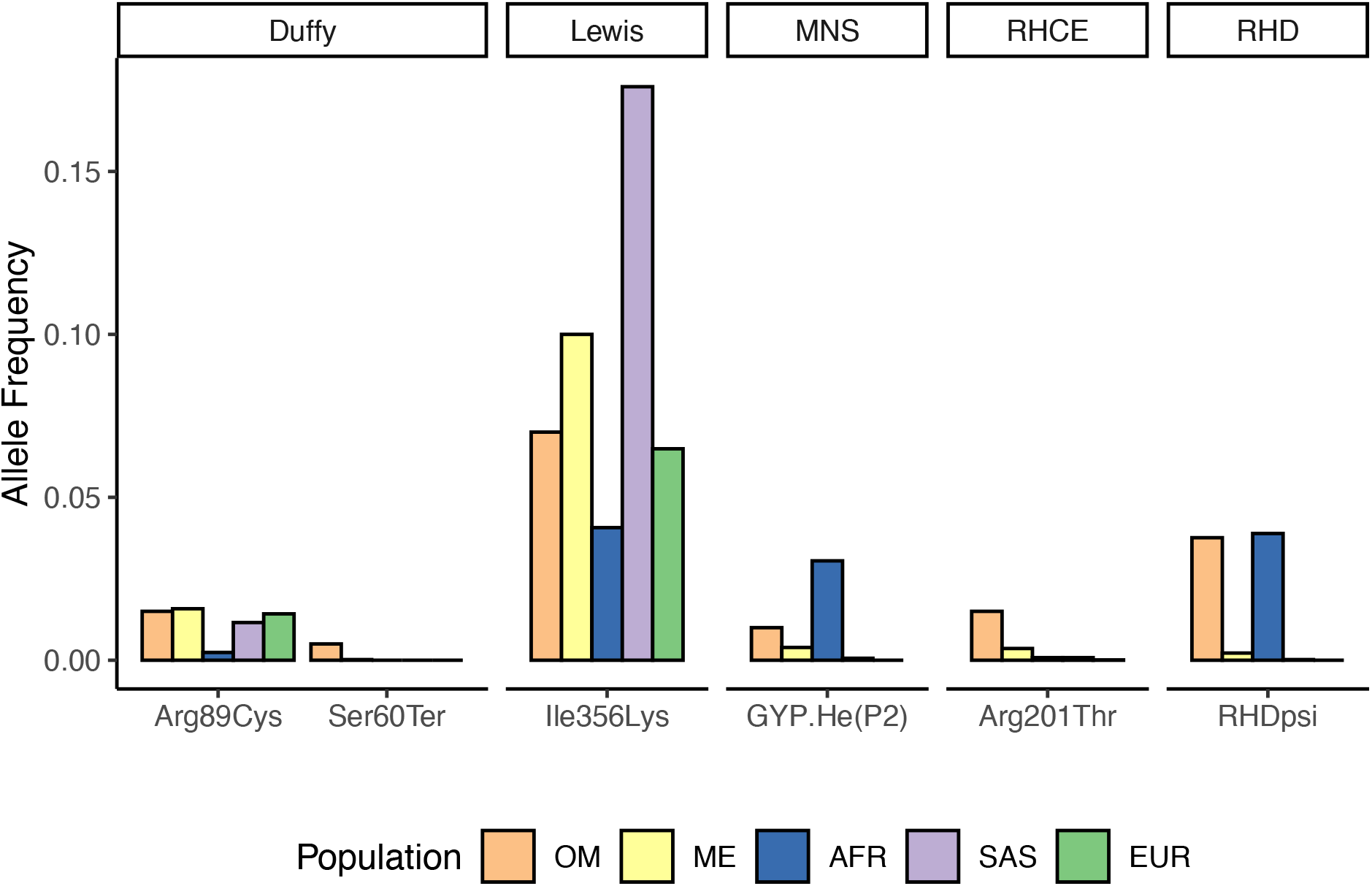
Allele frequencies of known blood group variants newly described in the Omani population (OM) compared to frequencies reported for Middle Eastern (ME), African (AFR), South Asian (SAS), and European (EUR) populations in gnomAD v4.0.

The E and e antigens, determined by alternative alleles at rs609320 (Ala226Pro), had 99% concordance. The discordant sample, inferred as E-e+ but serologically reported as E+e+, was found to carry an allele known to cause weak E expression, rs141398055 (Arg201Thr)^30^. Arg201Thr has the highest allele frequency in Middle Eastern populations (AF = 0.0036) in gnomADv4.0 and a frequency of 0.015 in the Omanis (Figure 1, Table 2).

The C antigen results from the presence of *RHD* exon 2 sequence in the paralogous location in *RHCE* (likely due to gene conversion), which can be detected as misalignment of reads to *RHD* exon 2. We initially used the approach implemented by Lane et al.^13^ comparing *RHCE* exon 2 coverage to the coverage of the *RHCE* locus. This resulted in a concordance of 91%. However, since the reads should be misaligning to *RHD* exon 2, we then compared the coverage across exon 2 of both genes in individuals that were hemizygous or homozygous for *RHD*. To account for hemizygosity, we adjusted the coverage range to > 0.67 for C-c+, 0.1-0.66 for C+c+, and < 0.1 for C+c-. Using this approach, the C and c antigens had 99% concordance. The discordant sample was inferred as C-c+ but serologically reported as C+c+ (Table S2) and remains unresolved as they are D negative so we could not apply this second approach.

### Duffy Blood Group

The genotype-phenotype concordance and resolving variants for the Duffy blood group in this dataset have been previously published in an analysis of genetic ancestry and positive selection at the Duffy blood group locus, *ACKR*1^19^. Briefly, inference of Fy^a^ and Fy^b^ antigen expression using rs12075 (Gly42Asp) and rs2814778 for the erythrocyte silent (ES) allele, Fy^ES^, resulted in a concordance of 96%. We found three discordant individuals genetically inferred as Fy(a-b+) but serologically reported as Fy(a-b-) to carry the Duffy X allele, Fy^X^ (rs34599082 Arg89Cys), that results in weak Fy^b^ expression and had previously not been described in Oman^31^. The fourth discordant individual was also genetically inferred as Fy(a-b+) and serologically reported as Fy(a-b-), but they did not carry the Fy^X^ allele. Instead, we identified a two base pair frameshift resulting in early protein termination (rs773692057 Ser62fs) carried by this individual, together with the Fy^ES^ allele causing the Duffy negative phenotype. The frameshift allele is rare but present in additional individuals from Oman and other populations in the Arabian Peninsula^19^.

### Lewis Blood Group

Two loci must be considered when inferring phenotypes of the Lewis blood group. The secretor, Le(a-b+), and non-secretor, Le(a+b-), phenotypes are most commonly determined by a nonsense mutation (rs601338 Trp154Ter) in *FUT2* on chromosome 19^32^ whereas the null phenotype, Le(a-b-), is caused by a variety of different alleles in *FUT3* that encode nonfunctional transferases, regardless of *FUT2* genotype^33^. We inferred the secretor phenotype using Trp154Ter in *FUT2* and the null phenotype using three different SNVs in *FUT3* that have been associated with Le(a-b-) in an Iranian population (rs28362459 Leu20Arg, rs812936 Arg68Trp, and rs778986 Met105Thr)^33^. This resulted in a concordance of 96%. When expanding to consider additional SNVs known to cause the null phenotype, we found that thirteen Omani individuals carried rs3894326 (Ile356Lys), a SNV commonly used for genotyping *FUT3* in European, South Asian and East Asian populations^34,35^ (Figure 1) leading to a revised concordance of 99%. The remaining discordant sample was inferred as Le(a-b+), but the serology reported them as Le(a+b+), a rare phenotype indicating a functional *FUT3* allele but a weak secretor allele at *FUT2*. This individual carried a unique missense variant, rs373779096 (Ala335Thr), that is very rare in gnomadv4.0 but primarily found in individuals of African/African American or admixed American ancestry (AF = 0.00037 and 0.00025 respectively) and in a single Middle Eastern individual (AF = 0.00017). Ala335Thr is located in the same exon as two other known weak secretor alleles (rs1047781 Ile140Phe and rs532253708 Met99Leu) that reduce enzymatic activity of alpha(1,2)fucosyltransferase^36-38^, and therefore may represent a new weak secretor allele, though further confirmation is needed.

### Lutheran Blood Group

The Lutheran blood group is encoded by the *BCAM* locus on chromosome 19. Using rs28399653 (Arg77His) to infer expression of Lu^a^ and Lu^b^ antigens, there was 99% concordance. The discordant sample was inferred as Lu(a-b+) by genotype but serologically reported as Lu(a-b-). The presence of Lu(a-b-) is consistent with the frequency observed in the previous study in Oman^12^, and suggests a higher frequency than elsewhere^39-45^. The Lu(a-b-) phenotype can either be due to homozygosity for loss of function alleles or expression of the *BCAM* gene below the level of detection by serology. The latter, referred to as In(Lu), is more common and has been attributed to heterozygous mutations affecting the transcription factor EKLF^44^. We looked for additional variants within the *BCAM* locus, in *EKLF*, and seven other erythroid transcription factors using the UCSC Genome Browser and JASPAR transcription factors tract^44,46-48^. We did not identify any loss-of-function alleles carried by this individual in *BCAM* or in *EKLF*. However, we did identify two adjacent SNVs falling in a GATA1 binding site for *SPI1* that are unique to this individual and thus a candidate for a new allele encoding a In(Lu) phenotype (rs533045163 and rs184739796). Neither SNV are reported in Middle Eastern individuals from the gnomADv4.0 database. Although rare, they are most common to African/African American individuals (AF = 0.006 for both SNVs).

### MNS Blood Group

The MNS blood group consists of 48 antigens encoded by three glycophorin genes on chromosome 4: *GYPA, GYPB*, and *GYPE*. For this analysis, we only analyzed the M, N, S, and s antigens. We inferred phenotypes using both SNVs and copy number calling as structural variants have been found to affect the glycophorin loci^1,25^. Expression of the M and N antigens can be inferred by three SNVs in *GYPA*: rs7682260 (Leu20Ser), and rs7687256 and rs7658293 (Glu24Gly)^49^. The resulting concordance was 94% (Figure 2A). Five of the six discordant samples were inferred as M-N+ whereas the remaining sample was M+N-. All six were reported as expressing both antigens (M+N+) by serology. It is likely the discordances are caused by difficulty aligning to this region due to the presence of the N antigen sequence in the reference at *GYPA* and *GYPB* and the M antigen sequence present in the reference at *GYPE*^*13*^. Additionally, we did not identify any structural variants that could be causing these discordances.

**Figure 2.**
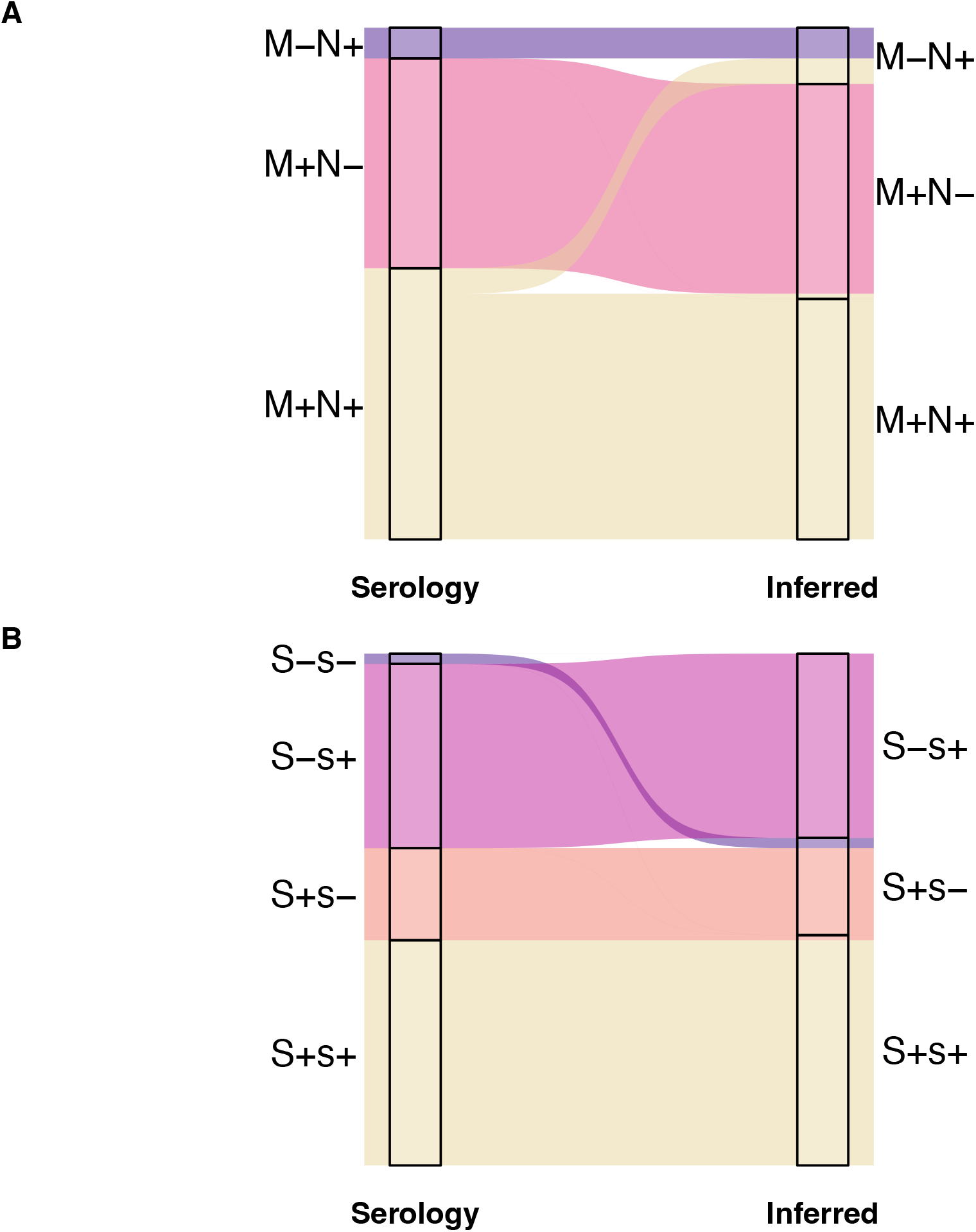
Alluvial plots showing the discordances in the MNS blood group. The left side of the plots are phenotypes determined by serology, and the right side are the phenotypes inferred by whole genome sequence data. The colors correspond to phenotypes inferred by serology. A) 88% concordance for the M and N antigens. B) 97% concordance for the S and s antigens.

The S and s antigens, inferred using rs7683365 (Thr48Met) in *GYPB*, had a concordance of 97% (Figure 2B). One discordant sample was inferred as S+s-but serologically found to be S-s-. We identified SNVs at multiple sites in *GYPB*, consistent with the Henshaw variant GYP.He(P2), a common cause of the S-s-U+^var^ phenotype in individuals with African ancestry^50^ (Figure 1). In addition to affecting 3 amino acids (Leu20Trp, Thr23Ser, Glu24Gly), the GYP.He(P2) variant includes another SNV on the same haplotype that alters a donor splice site in intron 5 (270+5 G>T, rs139511876) causing exon 5 to be skipped post-transcriptionally^50^. Copy number calling also revealed the presence of a *GYPB* gene deletion in this sample (Figure 3). Thus, the combination of GYP.He(P2) and the *GYPB* deletion is consistent with the S-s-phenotype.

**Figure 3.**
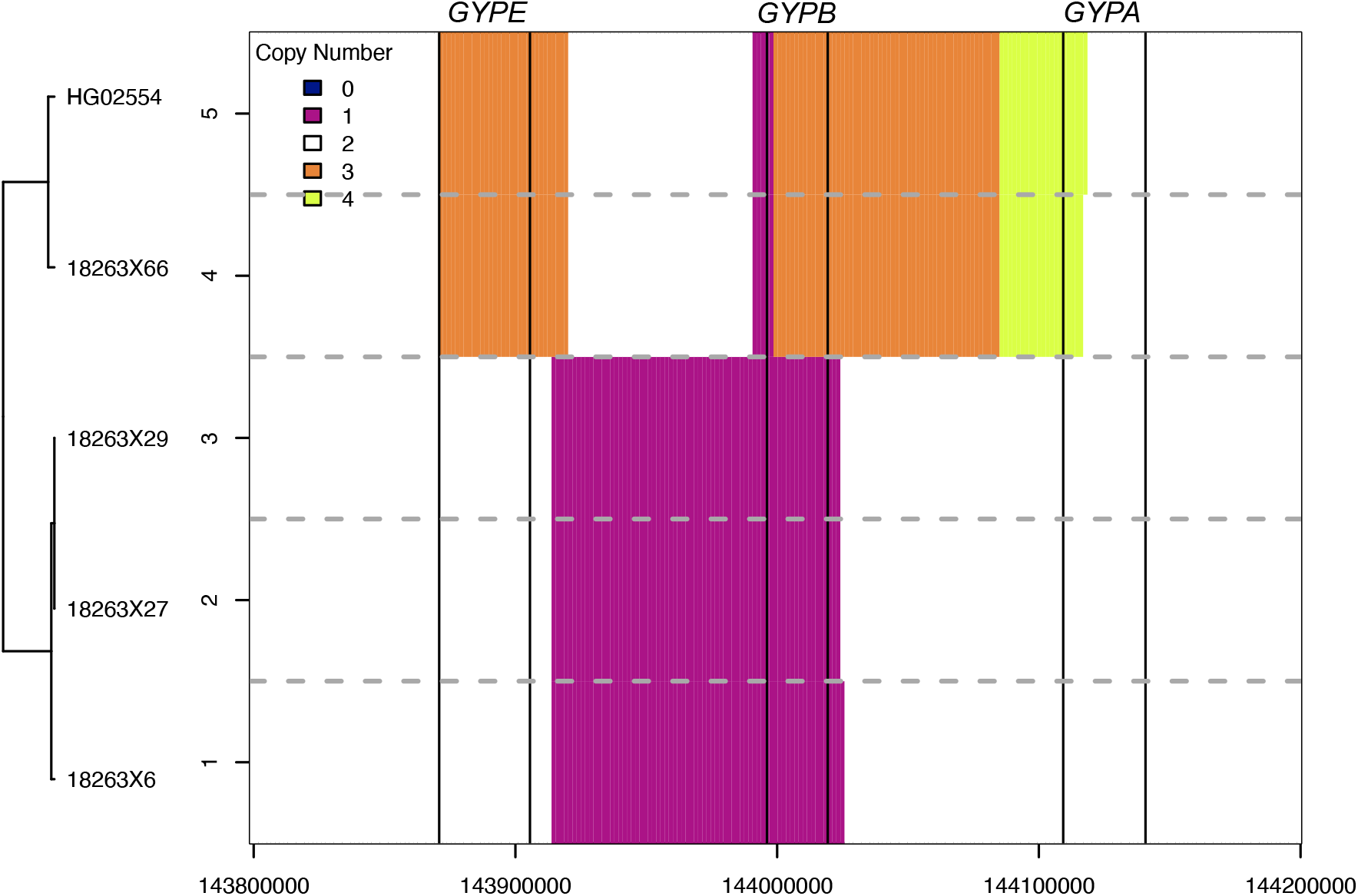
Copy number inference across the glycophorin genes based on read coverage. The x-axis corresponds to positions across the glycophorin gene region on chromosome 4. The vertical black bars indicate the genes from left to right: *GYPE, GYPB, GYPA*. The y-axis is labelled by each sample in the Omani dataset inferred as having a structural variant, except for HG02554 which is a 1000 genomes sample known to carry the Dantu structural variant. The dotted horizontal gray lines separate the inference for each sample. The colors correspond to the number of gene copies with white regions indicating two copies (no copy number variation relative to the reference).

The second discordant sample, inferred as S+s+ but serologically found to be S+s-, was inferred to carry the Dantu structural variant (previously identified in Haffener et al. 2024^19^). The Dantu allele has been associated with protection from severe *Plasmodium falciparum* malaria and is most common in East African populations^25^. However, the Dantu allele is thought to express s antigens^51^, so it is unclear if Dantu alone resolves this discordance. The remaining discordant sample was inferred as S+s-but serologically found to express both antigens (S+s+). This sample does not carry any known or novel missense, nonsense, or structural variants that we could identify.

### P1 Blood Group

We inferred the P_1_ and P_2_ phenotypes of the P1 blood group using four intronic variants in *A4GALT* on chromosome 22 reported to be associated with P1 antigen expression: rs66781836, rs5751348, rs8138197, and rs2143918^52^, although the causal variant remains unknown^53^. Phenotypes inferred with rs66781836 had the lowest accuracy with 97% concordance. The other three almost always occurred together and had a concordance of 99% suggesting rs66781836 is less likely to be the causal variant affecting P_1_ antigen expression, consistent with previous results^54,55^. The discordant sample, inferred as P_1_, was heterozygous for all three SNVs despite the serology reporting them as P_2_. Given that one of the three most likely causal variants, rs5751348, falls within an intronic transcription factor binding site^55^, we investigated other transcription factor binding sites within the *A4GALT* locus. We identified one potentially causal variant that falls within a STAT1 transcription factor binding site. This variant at position 22:42,721,266 is absent from gnomADv4.0 and dbSNP, suggesting it is extremely rare. However, two Omanis were found to be heterozygous: the discordant sample and a concordant sample homozygous for all three reference alleles serologically reported as P1.

## Discussion

Here, we comprehensively document the alleles underlying common blood group antigens in an Omani population by comparison of whole genome sequencing to serology. We demonstrate high concordance for all commonly tested blood group antigens in routine transfusion practice and report several recognized alleles altering blood group antigen expression that have previously not been described in Omanis. Notably, we describe alleles resolving discordances in the Rh, Duffy, Lewis, and MNS blood group systems (Table 2) in three or more unrelated individuals, suggesting these alleles are relatively common in the Omani population. Thus, these alleles should be considered for inclusion in red cell genotyping methods used in blood banks in Oman. Additionally, although singletons in this dataset, GYP.He(P2) and the 2 base pair *ACKR1* frameshift (Ser62fs), identified in a previous study with these samples^19^, should also be included given their resulting null phenotypes. These two alleles were also observed in other Arabian Peninsula and greater Middle Eastern populations indicating they are likely present throughout the region, although singletons in this dataset (Figure 1)^19^. Inclusion of additional populations from this region would provide a better understanding of how prevalent these alleles are and their relevance to red cell genotyping methods in other Arabian Peninsula populations.

While the overall blood group concordance was high and we were able to resolve discrepancies with this approach (revised overall concordance = 98.8%, Table 4), this analysis also revealed limitations to inferring MNS, ABO and Rh blood group phenotypes from whole genome sequencing data. The whole genome sequencing data had an average coverage of 16X and read length of 150 bp which we found led to mismapping in the glycophorin gene region and difficulty inferring M/N antigen expression as previously suggested^13^. An instance of the *ABO* O allele (Thr87AspfsTer107) being miscalled as a homozygote rather than a heterozygote also suggests a deeper coverage could improve blood group inferences using commonly typed SNVs. Lastly, the most unresolved discordances are in the MNS and Rh blood groups, likely given the complexity of these loci. Long-read sequencing or alignment to a pangenome, which can improve identification of structural variants^56,57^, may be necessary for reliable inference of these blood groups from DNA sequence.

**Table 4.** Adjusted per blood group concordances when including variants that resolved discrepancies (Table 2) between inferred phenotypes from whole genome sequence data and phenotypes reported by serology in the Omani samples.

| Blood Group | Average Concordance | Revised Concordance |
| --- | --- | --- |
| ABO | 98% | 100% |
| RHD | 99% | 100% |
| RHCE | 91% | 99% |
| Kell | 100% | 100% |
| Kidd | 100% | 100% |
| Duffy | 96% | 100% |
| Lewis | 96% | 99% |
| Lutheran | 99% | 99% |
| MNS | 91% | 92% |
| P1 | 99% | 99% |

Although discordant samples were not fully resolved in the MNS, Lewis, Lutheran, and P1 blood groups, we identified putatively causal variants that warrant further investigation. The Dantu SV identified in one of the discordant MNS samples has previously been believed to express the s+ antigen^51^. However, we did not identify any other potential causal variants within *GYPB* exons or cis-regulatory element binding sites that could have caused the S+s-phenotype in this individual. Within the Lewis blood group, we identified a SNV that could result in the weak secretor phenotype. Presently, only two weak secretor alleles have been described (Ile140Phe and Met99Leu)^36-38^ which reduce enzymatic activity and fall within the same exon of *FUT2* as the allele we identified, Ala335Thr. We also identified a candidate regulatory variant in a GATA1 transcription factor binding site of the erythrocyte-specific transcription factor locus, *SPI1* that could cause the In(Lu) phenotype as a result of reduced transcription of SPI1^58^, akin to the associated SNVs in a GATA1 binding site of *EKLF*^44^. Similarly, a candidate causal allele that falls within a binding site for the erythrocyte-specific transcription factor STAT1 in the 5’UTR of *A4GALT* may result in the absence of P1 antigen expression.

In conclusion, our findings document the alleles underlying common blood group antigens in Omanis and highlight the importance of considering population history when evaluating variants for blood group genotyping panels. For instance, the *RHD*\*ψ indel, erythrocyte-specific null allele in the Duffy blood group, and GYP.He(P2) alleles are most common to African populations. These alleles were found in multiple Omani blood donors from this dataset and their prevalence in the population is consistent with what is known about the shared genetic ancestry of the Omanis with East African populations^19,59,60^. Overall, this study emphasizes the necessity to increase population representation in genotype databases and indicates that whole genome sequencing paired with serology is a valuable approach for doing so in transfusion practice.

## Acknowledgments

This work was supported by the Omani Ministry of Higher Education, Innovation and Research (formally known as the Research Council, grant number RC/RG-MED/HAEM/18/02) and by the United States National Institutes of Health (grant number R35 GM147709 to E.M.L.). The support and resources from the Center for High Performance Computing at the University of Utah are gratefully acknowledged. The computational resources used were partially funded by the NIH Shared Instrumentation Grant 1S10OD021644-01A1. P.E.H. was supported by T32 GM007464 and T32 GM141848. Lastly, we thank all blood donors for participation in this study.

## Authorship Contributions

Conceptualization – A.Z.A. and E.M.L.; Methodology – P.E.H., S.A., A.A.S., and E.M.L.; Validation – E.M.L.; Formal Analysis – P.E.H.; Investigation – P.E.H., M.A., and S.A.H.; Resources – A.Z.A., and S.A.; Writing – Original Draft, P.E.H., A.Z.A., and E.M.L.; Writing – Review & Editing, A.Z.A. and S.A.; Visualization – P.E.H.; Supervision – A.Z.A., S.A., A.A.M., and E.M.L.; Project Administration – A.Z.A.; Funding Acquisition – A.Z.A. and E.M.L. All authors have read and approved this manuscript.

## References

1. ISBT. Red Cell Immunogenetics and Blood Group Terminology. Accessed February, 2024. https://www.isbtweb.org/isbt-working-parties/rcibgt.html

2. Genomes Project Consortium, Abecasis GR, Altshuler D, et al. A map of human genome variation from population-scale sequencing. Nature. Oct 28 2010;467(7319):1061–73. doi:10.1038/nature09534

3. Moller M, Joud M, Storry JR, Olsson ML. Erythrogene: a database for in-depth analysis of the extensive variation in 36 blood group systems in the 1000 Genomes Project. Blood Adv. Dec 27 2016;1(3):240–249. doi:10.1182/bloodadvances.2016001867

4. Montemayor-Garcia C, Karagianni P, Stiles DA, et al. Genomic coordinates and continental distribution of 120 blood group variants reported by the 1000 Genomes Project. Transfusion. Nov 2018;58(11):2693–2704. doi:10.1111/trf.14953

5. Al-Riyami AZ, Al Hinai D, Al-Rawahi M, et al. Molecular blood group screening in Omani blood donors. Vox Sang. Mar 2022;117(3):424–430. doi:10.1111/vox.13204

6. Ameen R, Al Shemmari S, Harris S, Teramura G, Delaney M. Classification of major and minor blood group antigens in the Kuwaiti Arab population. Transfusion and Apheresis Science. 2020/08/01/ 2020;59(4):102748. doi:10.1016/j.transci.2020.102748

7. El-Zawahri MM, Luqmani YA. Molecular genotyping and frequencies of A1, A2, B, O1 and O2 alleles of the ABO blood group system in a Kuwaiti population. Int J Hematol. Apr 2008;87(3):303–9. doi:10.1007/s12185-008-0036-0

8. Belali TM. Distribution of ABO and Rhesus Types in the Northern Asir Region in Saudi Arabia. J Blood Med. 2022;13:643–648. doi:10.2147/jbm.S383151

9. Bashwari LA, Al-Mulhim AA, Ahmad MS, Ahmed MA. Frequency of ABO blood groups in the Eastern region of Saudi Arabia. Saudi Med J. Nov 2001;22(11):1008–12.

10. Alzahrani M, Jawdat D, Alaskar A, Cereb N, Hajeer AH. ABO and Rh blood group genotypes in a cohort of Saudi stem cell donors. International Journal of Immunogenetics. 2018;45(2):63–64. doi:10.1111/iji.12354

11. Owaidah AY, Naffaa NM, Alumran A, Alzahrani F. Phenotype Frequencies of Major Blood Group Systems (Rh, Kell, Kidd, Duffy, MNS, P, Lewis, and Lutheran) Among Blood Donors in the Eastern Region of Saudi Arabia. J Blood Med. 2020;11:59–65. doi:10.2147/JBM.S236834

12. Al-Riyami AZ, Al-Marhoobi A, Al-Hosni S, et al. Prevalence of Red Blood Cell Major Blood Group Antigens and Phenotypes among Omani Blood Donors. Oman Med J. Nov 2019;34(6):496–503. doi:10.5001/omj.2019.92

13. Lane WJ, Westhoff CM, Gleadall NS, et al. Automated typing of red blood cell and platelet antigens: a whole-genome sequencing study. Lancet Haematol. Jun 2018;5(6):e241–e251. doi:10.1016/S2352-3026(18)30053-X

14. Al-Riyami AZ, Al-Mahrooqi S, Al-Hinai S, Al-Hosni S, Al-Madhani A, Daar S. Transfusion therapy and alloimmunization in Thalassemia Intermedia: a 10 year experience at a tertiary care university hospital. Transfus Apher Sci. Aug 2014;51(1):42–6. doi:10.1016/j.transci.2014.04.009

15. Al-Riyami AZ, Al-Muqbali A, Al-Sudiri S, et al. Risks of red blood cell alloimmunization in transfusion-dependent beta-thalassemia in Oman: a 25-year experience of a university tertiary care reference center and a literature review. Transfusion. Apr 2018;58(4):871–878. doi:10.1111/trf.14508

16. Al-Riyami AZ, Daar S. Transfusion in Haemoglobinopathies: Review and recommendations for local blood banks and transfusion services in Oman. Sultan Qaboos Univ Med J. Feb 2018;18(1):e3–e12. doi:10.18295/squmj.2018.18.01.002

17. Liu Z, Liu M, Mercado T, Illoh O, Davey R. Extended blood group molecular typing and next-generation sequencing. Transfus Med Rev. Oct 2014;28(4):177–86. doi:10.1016/j.tmrv.2014.08.003

18. Lane WJ, Westhoff CM, Uy JM, et al. Comprehensive red blood cell and platelet antigen prediction from whole genome sequencing: proof of principle. Transfusion. Mar 2016;56(3):743–54. doi:10.1111/trf.13416

19. Haffener PE, Al-Riyami AZ, Al-Zadjali S, et al. Adaptive admixture at ACKR1 (the Duffy locus) may have shaped Plasmodium vivax prevalence in Oman. bioRxiv. 2024:2024.03.06.583766. doi:10.1101/2024.03.06.583766

20. Li H. Aligning sequence reads, clone sequences and assembly contigs with BWA-MEM. arXiv:13033997. 2013;doi:10.48550/arXiv.1303.3997

21. Freed DN, Aldana R, Weber JA, Edwards JS. The Sentieon Genomics Tools -A fast and accurate solution to variant calling from next-generation sequence data. bioRxiv. May 12, 2017 2017;doi:10.1101/115717

22. Van der Auwera GA, Carneiro MO, Hartl C, et al. From FastQ data to high confidence variant calls: the Genome Analysis Toolkit best practices pipeline. Curr Protoc Bioinformatics. 2013;43(1110):11 10 1-11 10 33. doi:10.1002/0471250953.bi1110s43

23. Loh PR, Danecek P, Palamara PF, et al. Reference-based phasing using the Haplotype Reference Consortium panel. Nat Genet. Nov 2016;48(11):1443–1448. doi:10.1038/ng.3679

24. Li H, Handsaker B, Wysoker A, et al. The Sequence Alignment/Map format and SAMtools. Bioinformatics. Aug 15 2009;25(16):2078–9. doi:10.1093/bioinformatics/btp352

25. Leffler EM, Band G, Busby GBJ, et al. Resistance to malaria through structural variation of red blood cell invasion receptors. Science. Jun 16 2017;356(6343)doi:10.1126/science.aam6393

26. Yamamoto F, Clausen H, White T, Marken J, Hakomori S. Molecular genetic basis of the histo-blood group ABO system. Nature. May 17 1990;345(6272):229–33. doi:10.1038/345229a0

27. Wagner FF, Flegel WA. RHD gene deletion occurred in the Rhesus box. Blood. Jun 15 2000;95(12):3662–8.

28. Singleton BK, Green CA, Avent ND, et al. The presence of an RHD pseudogene containing a 37 base pair duplication and a nonsense mutation in africans with the Rh D-negative blood group phenotype. Blood. Jan 1 2000;95(1):12–8.

29. Chen S, Francioli LC, Goodrich JK, et al. A genome-wide mutational constraint map quantified from variation in 76,156 human genomes. bioRxiv. 2022:2022.03.20.485034. doi:10.1101/2022.03.20.485034

30. Bugert P, Scharberg EA, Geisen C, von Zabern I, Flegel WA. RhCE protein variants in Southwestern Germany detected by serologic routine testing. Transfusion. Sep 2009;49(9):1793–802. doi:10.1111/j.1537-2995.2009.02220.x

31. Tournamille C, Le Van Kim C, Gane P, et al. Arg89Cys substitution results in very low membrane expression of the Duffy antigen/receptor for chemokines in Fy(x) individuals. Blood. Sep 15 1998;92(6):2147–56.

32. Daniels G. The molecular genetics of blood group polymorphism. Hum Genet. Dec 2009;126(6):729–42. doi:10.1007/s00439-009-0738-2

33. Farahmand M, Jalilvand S, Arashkia A, et al. Estimation of genetic variation in the Secretor and Lewis genes in Iranian hospitalized children. Transfus Clin Biol. Feb 2021;28(1):11–15. doi:10.1016/j.tracli.2020.12.001

34. Hu D, Zhang D, Zheng S, Guo M, Lin X, Jiang Y. Association of Ulcerative Colitis with FUT2 and FUT3 Polymorphisms in Patients from Southeast China. PLoS One. 2016;11(1):e0146557. doi:10.1371/journal.pone.0146557

35. Taylor-Cousar JL, Zariwala MA, Burch LH, et al. Histo-blood group gene polymorphisms as potential genetic modifiers of infection and cystic fibrosis lung disease severity. PLoS One. 2009;4(1):e4270. doi:10.1371/journal.pone.0004270

36. Soejima M, Koda Y. Detection of the weak-secretor rs1047781 (385A>T) single nucleotide polymorphism using an unlabeled probe high-resolution melting-based method. ELECTROPHORESIS. 2021;42(12-13):1362-1365. doi:10.1002/elps.202000386

37. Soejima M, Koda Y. Survey and characterization of nonfunctional alleles of FUT2 in a database. Sci Rep. Feb 4 2021;11(1):3186. doi:10.1038/s41598-021-82895-w

38. Henry S, Mollicone R, Fernandez P, Samuelsson B, Oriol R, Larson G. Molecular basis for erythrocyte Le(a+ b+) and salivary ABH partial-secretor phenotypes: expression of a FUT2 secretor allele with an A-->T mutation at nucleotide 385 correlates with reduced alpha(1,2) fucosyltransferase activity. Glycoconj J. Dec 1996;13(6):985–93. doi:10.1007/bf01053194

39. Rowe GP, Gale SA, Daniels GL, Green CA, Tippett P. A study on Lu-null families in South Wales. Ann Hum Genet. Jul 1992;56(3):267–72. doi:10.1111/j.1469-1809.1992.tb01151.x

40. Darnborough J, Firth R, Biles CM, Goldsmith KL, Crawford MN. A “new” antibody anti-Lu-a-Lu-b and two further examples of the genotype Lu(a-b-). Nature. May 25 1963;198:796. doi:10.1038/198796a0

41. Brown F, Simpson S, Cornwall S, Moore BP, Oyen R, Marsh WL. The recessive Lu(a-b-) phenotype. A family study. Vox Sang. Mar 1974;26(3):259–64. doi:10.1111/j.1423-0410.1974.tb02694.x

42. Myhre B, Thompson M, Anson C, Fishkin B, Carter PK. A Further Example of the Recessive Lu(a-b-) Phenotype. Vox Sanguinis. 1975;29(1):66–68. doi:10.1111/j.1423-0410.1975.tb00478.x

43. Karamatic Crew V, Mallinson G, Green C, et al. Different inactivating mutations in the LU genes of three individuals with the Lutheran-null phenotype. Transfusion. Mar 2007;47(3):492–8. doi:10.1111/j.1537-2995.2006.01141.x

44. Singleton BK, Burton NM, Green C, Brady RL, Anstee DJ. Mutations in EKLF/KLF1 form the molecular basis of the rare blood group In(Lu) phenotype. Blood. Sep 1 2008;112(5):2081–8. doi:10.1182/blood-2008-03-145672

45. Singleton BK, Roxby DJ, Stirling JW, et al. A novel GATA1 mutation (Stop414Arg) in a family with the rare X-linked blood group Lu(a-b-) phenotype and mild macrothrombocytic thrombocytopenia. Br J Haematol. Apr 2013;161(1):139–42. doi:10.1111/bjh.12184

46. Kent WJ, Sugnet CW, Furey TS, et al. The human genome browser at UCSC. Genome Res. Jun 2002;12(6):996–1006. doi:10.1101/gr.229102

47. Castro-Mondragon JA, Riudavets-Puig R, Rauluseviciute I, et al. JASPAR 2022: the 9th release of the open-access database of transcription factor binding profiles. Nucleic Acids Res. Jan 7 2022;50(D1):D165-D173. doi:10.1093/nar/gkab1113

48. Rauluseviciute I, Riudavets-Puig R, Blanc-Mathieu R, et al. JASPAR 2024: 20th anniversary of the open-access database of transcription factor binding profiles. Nucleic Acids Res. Jan 5 2024;52(D1):D174-D182. doi:10.1093/nar/gkad1059

49. Furst D, Tsamadou C, Neuchel C, Schrezenmeier H, Mytilineos J, Weinstock C. Next-Generation Sequencing Technologies in Blood Group Typing. Transfus Med Hemother. Feb 2020;47(1):4–13. doi:10.1159/000504765

50. Storry JR, Reid ME, Fetics S, Huang CH. Mutations in GYPB exon 5 drive the S-s-U+(var) phenotype in persons of African descent: implications for transfusion. Transfusion. Dec 2003;43(12):1738–47. doi:10.1046/j.0041-1132.2003.00585.x

51. Blumenfeld OO, Smith AJ, Moulds JJ. Membrane glycophorins of Dantu blood group erythrocytes. J Biol Chem. Aug 25 1987;262(24):11864–70.

52. Lane WJ, Aguad M, Smeland-Wagman R, et al. A whole genome approach for discovering the genetic basis of blood group antigens: independent confirmation for P1 and Xg(a). Transfusion. Mar 2019;59(3):908–915. doi:10.1111/trf.15089

53. Stenfelt L, Hellberg A, Westman JS, Olsson ML. The P1PK blood group system: revisited and resolved. Immunohematology. Sep 2020;36(3):99–103.

54. Lai YJ, Wu WY, Yang CM, et al. A systematic study of single-nucleotide polymorphisms in the A4GALT gene suggests a molecular genetic basis for the P1/P2 blood groups. Transfusion. Dec 2014;54(12):3222–31. doi:10.1111/trf.12771

55. Westman JS, Stenfelt L, Vidovic K, et al. Allele-selective RUNX1 binding regulates P1 blood group status by transcriptional control of A4GALT. Blood. Apr 5 2018;131(14):1611–1616. doi:10.1182/blood-2017-08-803080

56. Liao WW, Asri M, Ebler J, et al. A draft human pangenome reference. Nature. May 2023;617(7960):312–324. doi:10.1038/s41586-023-05896-x

57. Miller DE, Sulovari A, Wang T, et al. Targeted long-read sequencing identifies missing disease-causing variation. The American Journal of Human Genetics. 2021/08/05/ 2021;108(8):1436–1449. doi:10.1016/j.ajhg.2021.06.006

58. Ballas SK, Marcolina MJ, Crawford MN. In vitro storage and in vivo survival studies of red cells from persons with the In(Lu) gene. Transfusion. Sep 1992;32(7):607–11. doi:10.1046/j.1537-2995.1992.32792391031.x

59. Fernandes V, Brucato N, Ferreira JC, et al. Genome-Wide Characterization of Arabian Peninsula Populations: Shedding Light on the History of a Fundamental Bridge between Continents. Mol Biol Evol. Mar 1 2019;36(3):575–586. doi:10.1093/molbev/msz005

60. Ferreira JC, Alshamali F, Montinaro F, et al. Projecting Ancient Ancestry in Modern-Day Arabians and Iranians: A Key Role of the Past Exposed Arabo-Persian Gulf on Human Migrations. Genome Biol Evol. Sep 1 2021;13(9)doi:10.1093/gbe/evab194

