## Supplemental Tables for "Characterization of Blood Group Variants in an Omani Population by Comparison of Whole Genome Sequencing and Serology"

**Table S1**. Graded serology (0-4) for ABO, RHD, RHCE, Kell, Kidd, Duffy,  Lewis, Lutheran, MNS, and P1, blood group systems for all 100 Omani blood donors.

| Research sequence# | Blood group | Rh+K | | | | P | Lewis | | Lutheran | | Kell | | | | Kid | | MNS | | | | Duffy | |
|---|---|---|---|---|---|---|---|---|---|---|---|---|---|---|---|---|---|---|---|---|---|---|
| | | C | c | E | e | P1 | Le(a) | Le(b) | Lu(a) | Lu(b) | K | k | Kp(a) | Kp(b) | Jk(a) | Jk(b) | M | N | S | s | Fy(a) | Fy(b) |
| 18263X1 | O Rh+ | 0 | 4 | 4 | 4 | 0 | 0 | 4 | 0 | 3 | 4 | 3 | 0 | 3 | 0 | 4 | 4 | 4 | 3 | 3 | 0 | 0 |
| 18263X2 | O Rh+ | 4 | 4 | 4 | 4 | 4 | 4 | 0 | 0 | 3 | 0 | 3 | 0 | 3 | 0 | 4 | 4 | 4 | 3 | 3 | 0 | 0 |
| 18263X3 | O Rh+ | 0 | 4 | 4 | 4 | 0 | 0 | 4 | 0 | 3 | 0 | 3 | 3 | 3 | 0 | 4 | 0 | 4 | 0 | 3 | 0 | 0 |
| 18263X4 | A Rh+ | 0 | 4 | 4 | 4 | 4 | 0 | 2 | 0 | 3 | 0 | 3 | 0 | 3 | 4 | 0 | 4 | 4 | 3 | 3 | 0 | 0 |
| 18263X5 | A Rh+ | 4 | 4 | 4 | 4 | 4 | 0 | 4 | 0 | 3 | 0 | 4 | 0 | 3 | 4 | 0 | 4 | 4 | 3 | 3 | 0 | 0 |
| 18263X6 | O Rh+ | 4 | 4 | 4 | 4 | 4 | 0 | 4 | 0 | 2 | 0 | 3 | 0 | 3 | 4 | 4 | 4 | 4 | 1 | 0 | 0 | 0 |
| 18263X7 | O Rh+ | 4 | 4 | 0 | 4 | 4 | 0 | 4 | 0 | 2 | 0 | 3 | 0 | 3 | 4 | 0 | 4 | 0 | 3 | 3 | 0 | 0 |
| 18263X8 | A Rh+ | 4 | 4 | 0 | 4 | 0 | 0 | 4 | 0 | 3 | 0 | 3 | 0 | 3 | 4 | 0 | 4 | 4 | 0 | 3 | 0 | 0 |
| 18263X9 | A Rh+ | 4 | 4 | 0 | 4 | 4 | 3 | 4 | 0 | 1 | 0 | 3 | 0 | 3 | 4 | 4 | 4 | 0 | 0 | 3 | 0 | 0 |
| 18263X10 | A Rh+ | 4 | 4 | 4 | 4 | 0 | 0 | 4 | 0 | 3 | 0 | 3 | 0 | 3 | 4 | 0 | 4 | 4 | 3 | 3 | 0 | 0 |
| 18263X11 | O Rh+ | 4 | 0 | 0 | 4 | 4 | 0 | 0 | 0 | 1 | 0 | 3 | 0 | 3 | 0 | 4 | 4 | 0 | 3 | 0 | 0 | 3 |
| 18263X12 | A Rh- | 0 | 4 | 0 | 4 | 4 | 0 | 4 | 0 | 3 | 0 | 3 | 0 | 3 | 4 | 0 | 4 | 4 | 0 | 3 | 0 | 0 |
| 18263X13 | O Rh+ | 4 | 4 | 0 | 4 | 4 | 0 | 0 | 0 | 3 | 0 | 3 | 0 | 3 | 4 | 0 | 4 | 4 | 0 | 3 | 0 | 0 |
| 18263X14 | A Rh+ | 4 | 0 | 0 | 4 | 4 | 0 | 0 | 0 | 3 | 0 | 3 | 0 | 3 | 4 | 0 | 4 | 0 | 3 | 0 | 0 | 0 |
| 18263X15 | O Rh+ | 4 | 0 | 0 | 4 | 0 | 0 | 4 | 0 | 3 | 0 | 3 | 0 | 3 | 4 | 4 | 4 | 0 | 3 | 0 | 0 | 0 |
| 18263X16 | AB Rh+ | 0 | 4 | 0 | 4 | 4 | 0 | 3 | 0 | 3 | 0 | 3 | 0 | 3 | 0 | 4 | 4 | 4 | 3 | 3 | 3 | 0 |
| 18263X17 | O Rh+ | 4 | 4 | 0 | 4 | 4 | 0 | 0 | 0 | 3 | 0 | 3 | 0 | 3 | 4 | 4 | 3 | 4 | 0 | 3 | 0 | 3 |
| 18263X18 | O Rh+ | 4 | 0 | 0 | 4 | 3 | 0 | 0 | 0 | 3 | 0 | 3 | 0 | 3 | 0 | 4 | 4 | 0 | 3 | 0 | 0 | 0 |
| 18263X19 | O Rh- | 0 | 4 | 0 | 4 | 3 | 0 | 4 | 0 | 3 | 0 | 3 | 0 | 3 | 4 | 0 | 4 | 0 | 3 | 3 | 0 | 0 |
| 18263X20 | O Rh+ | 4 | 0 | 0 | 4 | 4 | 4 | 0 | 0 | 3 | 0 | 3 | 0 | 3 | 0 | 4 | 4 | 0 | 3 | 3 | 0 | 0 |
| 18263X21 | A Rh+ | 4 | 0 | 0 | 4 | 4 | 0 | 4 | 0 | 3 | 0 | 3 | 0 | 3 | 4 | 0 | 4 | 4 | 0 | 3 | 0 | 0 |
| 18263X22 | O Rh+ | 4 | 4 | 4 | 4 | 4 | 0 | 4 | 0 | 3 | 0 | 3 | 0 | 3 | 4 | 4 | 4 | 4 | 0 | 3 | 0 | 0 |
| 18263X23 | B Rh+ | 4 | 0 | 0 | 4 | 3 | 0 | 4 | 2 | 3 | 0 | 3 | 0 | 3 | 4 | 4 | 4 | 4 | 3 | 3 | 0 | 0 |
| 18263X24 | A Rh+ | 4 | 4 | 4 | 4 | 4 | 0 | 3 | 0 | 1 | 0 | 4 | 0 | 3 | 4 | 4 | 4 | 4 | 0 | 3 | 0 | 0 |
| 18263X25 | O Rh+ | 4 | 4 | 0 | 4 | 4 | 0 | 4 | 0 | 1 | 4 | 3 | 0 | 3 | 4 | 4 | 4 | 0 | 3 | 0 | 0 | 0 |
| 18263X26 | O Rh+ | 0 | 4 | 0 | 4 | 4 | 4 | 0 | 0 | 2 | 0 | 3 | 0 | 4 | 4 | 0 | 4 | 4 | 0 | 3 | 0 | 0 |
| 18263X27 | B Rh+ | 0 | 4 | 4 | 0 | 4 | 0 | 4 | 0 | 1 | 0 | 3 | 0 | 3 | 4 | 0 | 4 | 3 | 0 | 0 | 0 | 0 |
| 18263X28 | O Rh+ | 4 | 4 | 0 | 4 | 4 | 0 | 4 | 0 | 3 | 0 | 3 | 0 | 3 | 4 | 4 | 4 | 4 | 3 | 3 | 0 | 0 |
| 18263X29 | A Rh+ | 0 | 4 | 0 | 4 | 4 | 4 | 0 | 0 | 3 | 0 | 3 | 0 | 3 | 4 | 0 | 0 | 4 | 0 | 3 | 0 | 3 |
| 18263X30 | O Rh+ | 4 | 4 | 0 | 4 | 4 | 0 | 4 | 0 | 3 | 4 | 3 | 0 | 3 | 0 | 4 | 4 | 0 | 3 | 3 | 0 | 3 |

| | | | | | | | | | | | | | | | | | | | | | | |
|---|---|---|---|---|---|---|---|---|---|---|---|---|---|---|---|---|---|---|---|---|---|---|
| 18263X31 | O Rh+ | 4 | 4 | 0 | 4 | 4 | 0 | 4 | 0 | 3 | 0 | 3 | 0 | 3 | 4 | 0 | 4 | 4 | 3 | 3 | 0 | 0 |
| 18263X32 | O Rh+ | 4 | 0 | 0 | 4 | 0 | 3 | 0 | 0 | 3 | 0 | 3 | 0 | 3 | 4 | 0 | 4 | 1 | 3 | 0 | 0 | 0 |
| 18263X33 | AB Rh+ | 0 | 4 | 0 | 4 | 4 | 0 | 3 | 0 | 2 | 0 | 3 | 0 | 3 | 4 | 0 | 4 | 0 | 3 | 3 | 0 | 0 |
| 18263X34 | O Rh+ | 4 | 0 | 4 | 4 | 2 | 3 | 0 | 0 | 0 | 0 | 3 | 0 | 3 | 0 | 4 | 4 | 4 | 0 | 3 | 0 | 0 |
| 18263X35 | O Rh+ | 4 | 0 | 0 | 4 | 0 | 0 | 4 | 0 | 1 | 0 | 3 | 0 | 3 | 4 | 0 | 4 | 4 | 3 | 0 | 0 | 0 |
| 18263X36 | O Rh+ | 4 | 4 | 0 | 4 | 0 | 0 | 0 | 0 | 2 | 0 | 3 | 0 | 3 | 0 | 4 | 3 | 3 | 3 | 0 | 0 | 0 |
| 18263X37 | O Rh+ | 4 | 0 | 0 | 4 | 3 | 0 | 4 | 0 | 2 | 0 | 3 | 0 | 3 | 4 | 3 | 3 | 3 | 0 | 3 | 0 | 0 |
| 18263X38 | O Rh+ | 0 | 4 | 4 | 4 | 3 | 3 | 0 | 0 | 2 | 0 | 2 | 0 | 3 | 3 | 3 | 3 | 0 | 3 | 3 | 0 | 0 |
| 18263X39 | O Rh+ | 4 | 4 | 4 | 4 | 3 | 0 | 4 | 0 | 2 | 0 | 3 | 0 | 3 | 4 | 0 | 2 | 3 | 3 | 3 | 0 | 0 |
| 18263X40 | A Rh- | 0 | 4 | 0 | 4 | 3 | 0 | 3 | 0 | 2 | 0 | 3 | 0 | 2 | 3 | 0 | 4 | 0 | 3 | 3 | 3 | 0 |
| 18263X41 | O Rh+ | 4 | 0 | 0 | 4 | 0 | 0 | 0 | 0 | 2 | 0 | 3 | 0 | 3 | 4 | 0 | 3 | 0 | 3 | 3 | 0 | 0 |
| 18263X42 | O Rh+ | 0 | 4 | 0 | 4 | 2 | 0 | 4 | 0 | 2 | 0 | 3 | 0 | 3 | 3 | 3 | 3 | 0 | 0 | 3 | 0 | 0 |
| 18263X43 | A Rh+ | 4 | 0 | 0 | 4 | 3 | 0 | 4 | 0 | 2 | 0 | 3 | 0 | 3 | 4 | 0 | 3 | 0 | 2 | 0 | 0 | 0 |
| 18263X44 | O Rh+ | 4 | 4 | 0 | 4 | 3 | 0 | 4 | 0 | 2 | 0 | 3 | 0 | 3 | 3 | 3 | 3 | 0 | 2 | 2 | 0 | 0 |
| 18263X45 | A Rh+ | 4 | 0 | 0 | 4 | 3 | 0 | 3 | 0 | 2 | 0 | 3 | 0 | 3 | 4 | 0 | 3 | 0 | 2 | 2 | 3 | 2 |
| 18263X46 | AB Rh+ | 4 | 0 | 0 | 4 | 2 | 0 | 0 | 0 | 2 | 0 | 3 | 0 | 3 | 3 | 3 | 3 | 0 | 0 | 3 | 0 | 0 |
| 18263X47 | O Rh+ | 4 | 4 | 0 | 4 | 3 | 0 | 0 | 0 | 2 | 0 | 3 | 0 | 3 | 3 | 3 | 3 | 0 | 2 | 2 | 0 | 0 |
| 18263X48 | AB Rh+ | 4 | 0 | 0 | 4 | 3 | 0 | 0 | 0 | 2 | 0 | 3 | 0 | 3 | 3 | 0 | 4 | 0 | 3 | 0 | 0 | 0 |
| 18263X49 | O Rh+ | 4 | 4 | 0 | 4 | 2 | 0 | 3 | 0 | 2 | 0 | 3 | 0 | 3 | 4 | 3 | 3 | 3 | 2 | 2 | 0 | 0 |
| 18263X50 | O Rh+ | 4 | 0 | 0 | 4 | 3 | 0 | 0 | 0 | 2 | 0 | 2 | 0 | 3 | 4 | 3 | 2 | 3 | 2 | 0 | 0 | 0 |
| 18263X51 | A Rh- | 0 | 4 | 0 | 4 | 3 | 0 | 3 | 0 | 2 | 0 | 3 | 0 | 3 | 4 | 0 | 3 | 0 | 2 | 0 | 0 | 0 |
| 18263X52 | O Rh+ | 4 | 4 | 0 | 4 | 3 | 0 | 4 | 0 | 3 | 0 | 3 | 0 | 3 | 4 | 3 | 4 | 0 | 0 | 2 | 0 | 0 |
| 18263X53 | O Rh- | 0 | 4 | 0 | 4 | 0 | 2 | 0 | 0 | 2 | 0 | 2 | 0 | 3 | 4 | 0 | 4 | 0 | 3 | 2 | 0 | 0 |
| 18263X54 | O Rh+ | 4 | 0 | 0 | 4 | 4 | 0 | 0 | 0 | 3 | 0 | 3 | 0 | 3 | 4 | 3 | 4 | 0 | 3 | 0 | 0 | 0 |
| 18263X55 | O Rh+ | 4 | 0 | 0 | 4 | 0 | 3 | 0 | 0 | 2 | 0 | 3 | 0 | 3 | 3 | 4 | 4 | 0 | 0 | 3 | 0 | 0 |
| 18263X56 | B Rh+ | 4 | 4 | 0 | 4 | 3 | 2 | 0 | 0 | 2 | 0 | 3 | 0 | 3 | 4 | 3 | 3 | 3 | 0 | 3 | 0 | 0 |
| 18263X57 | O Rh+ | 0 | 4 | 0 | 4 | 3 | 0 | 3 | 0 | 2 | 4 | 3 | 0 | 2 | 4 | 0 | 3 | 3 | 3 | 3 | 0 | 0 |
| 18263X58 | O Rh+ | 4 | 0 | 0 | 4 | 0 | 0 | 3 | 0 | 2 | 0 | 3 | 3 | 2 | 4 | 0 | 2 | 2 | 2 | 2 | 2 | 0 |

| | | | | | | | | | | | | | | | | | | | | | | |
|---|---|---|---|---|---|---|---|---|---|---|---|---|---|---|---|---|---|---|---|---|---|---|
| 18263X59 | A Rh+ | 4 | 0 | 0 | 4 | 2 | 0 | 3 | 0 | 2 | 0 | 3 | 0 | 3 | 4 | 0 | 3 | 0 | 2 | 3 | 0 | 0 |
| 18263X60 | O Rh+ | 4 | 0 | 0 | 4 | 3 | 0 | 4 | 0 | 2 | 0 | 3 | 0 | 3 | 3 | 4 | 3 | 0 | 2 | 2 | 0 | 0 |
| 18263X61 | B Rh+ | 0 | 4 | 4 | 4 | 3 | 3 | 0 | 0 | 2 | 0 | 2 | 0 | 3 | 3 | 3 | 4 | 0 | 2 | 2 | 0 | 0 |
| 18263X62 | O Rh+ | 4 | 4 | 0 | 4 | 3 | 0 | 4 | 0 | 3 | 0 | 3 | 0 | 3 | 4 | 0 | 3 | 3 | 2 | 2 | 0 | 0 |
| 18263X63 | B Rh+ | 4 | 0 | 0 | 4 | 4 | 0 | 3 | 0 | 3 | 4 | 2 | 0 | 3 | 0 | 3 | 3 | 0 | 2 | 2 | 0 | 0 |
| 18263X64 | B Rh+ | 0 | 4 | 4 | 0 | 4 | 0 | 3 | 0 | 2 | 0 | 3 | 0 | 2 | 0 | 3 | 0 | 3 | 2 | 3 | 0 | 0 |
| 18263X65 | O Rh- | 0 | 4 | 0 | 4 | 3 | 3 | 0 | 0 | 2 | 0 | 3 | 0 | 2 | 4 | 0 | 2 | 3 | 3 | 2 | 0 | 0 |
| 18263X66 | A Rh- | 4 | 4 | 0 | 4 | 2 | 0 | 0 | 0 | 2 | 0 | 3 | 0 | 3 | 4 | 0 | 2 | 3 | 2 | 0 | 2 | 0 |
| 18263X67 | O Rh+ | 0 | 4 | 4 | 4 | 4 | 0 | 4 | 0 | 2 | 0 | 3 | 0 | 3 | 3 | 3 | 2 | 3 | 2 | 2 | 0 | 0 |
| 18263X68 | O Rh+ | 4 | 4 | 0 | 4 | 0 | 0 | 3 | 0 | 3 | 0 | 3 | 0 | 3 | 4 | 0 | 2 | 3 | 0 | 2 | 0 | 0 |
| 18263X69 | O Rh- | 0 | 4 | 0 | 4 | 2 | 0 | 3 | 0 | 2 | 0 | 3 | 0 | 3 | 4 | 4 | 3 | 0 | 0 | 3 | 0 | 2 |
| 18263X70 | O Rh+ | 4 | 4 | 4 | 4 | 0 | 0 | 4 | 0 | 2 | 4 | 2 | 0 | 3 | 4 | 0 | 3 | 0 | 2 | 3 | 0 | 0 |
| 18263X71 | O Rh+ | 4 | 4 | 0 | 4 | 3 | 0 | 0 | 0 | 2 | 0 | 3 | 0 | 3 | 4 | 0 | 2 | 3 | 2 | 2 | 0 | 0 |
| 18263X72 | B Rh+ | 0 | 4 | 4 | 4 | 0 | 0 | 3 | 0 | 2 | 0 | 3 | 0 | 3 | 3 | 0 | 2 | 3 | 0 | 2 | 0 | 2 |
| 18263X73 | A Rh+ | 4 | 0 | 0 | 4 | 3 | 0 | 4 | 0 | 2 | 0 | 3 | 0 | 3 | 4 | 0 | 3 | 0 | 2 | 3 | 2 | 0 |
| 18263X74 | O Rh+ | 4 | 4 | 0 | 4 | 3 | 0 | 4 | 0 | 2 | 0 | 3 | 0 | 3 | 0 | 4 | 3 | 3 | 3 | 3 | 0 | 0 |
| 18263X75 | B Rh+ | 0 | 4 | 4 | 4 | 0 | 0 | 4 | 0 | 2 | 0 | 3 | 0 | 2 | 4 | 4 | 3 | 3 | 3 | 3 | 0 | 0 |
| 18263X76 | B Rh+ | 4 | 4 | 0 | 4 | 2 | 0 | 3 | 0 | 2 | 0 | 3 | 0 | 3 | 4 | 0 | 3 | 3 | 0 | 3 | 0 | 0 |
| 18263X77 | O Rh+ | 4 | 4 | 0 | 4 | 3 | 0 | 0 | 0 | 3 | 0 | 3 | 0 | 2 | 3 | 3 | 2 | 3 | 2 | 3 | 0 | 0 |
| 18263X78 | B Rh+ | 4 | 4 | 4 | 4 | 4 | 0 | 3 | 0 | 2 | 0 | 3 | 0 | 3 | 0 | 3 | 4 | 0 | 0 | 3 | 0 | 0 |
| 18263X79 | O Rh+ | 4 | 4 | 0 | 4 | 3 | 0 | 4 | 0 | 2 | 0 | 3 | 0 | 3 | 0 | 4 | 3 | 3 | 0 | 3 | 0 | 0 |
| 18263X80 | O Rh+ | 4 | 0 | 0 | 4 | 2 | 3 | 0 | 0 | 2 | 0 | 3 | 0 | 3 | 4 | 3 | 3 | 3 | 2 | 2 | 0 | 0 |
| 18263X81 | O Rh+ | 4 | 0 | 0 | 4 | 3 | 0 | 4 | 0 | 2 | 0 | 3 | 0 | 3 | 4 | 0 | 3 | 3 | 3 | 3 | 2 | 0 |
| 18263X82 | O Rh+ | 0 | 4 | 4 | 4 | 3 | 0 | 4 | 0 | 2 | 0 | 3 | 0 | 2 | 4 | 0 | 2 | 2 | 0 | 2 | 0 | 0 |
| 18263X83 | O Rh+ | 0 | 4 | 0 | 4 | 4 | 0 | 4 | 0 | 2 | 0 | 3 | 0 | 3 | 4 | 4 | 4 | 0 | 3 | 0 | 0 | 0 |
| 18263X84 | O Rh+ | 4 | 4 | 0 | 4 | 4 | 0 | 4 | 0 | 2 | 0 | 3 | 0 | 3 | 4 | 0 | 2 | 2 | 0 | 2 | 0 | 0 |
| 18263X85 | A Rh+ | 0 | 4 | 0 | 4 | 2 | 0 | 4 | 0 | 2 | 0 | 2 | 0 | 3 | 4 | 0 | 3 | 2 | 0 | 3 | 0 | 0 |
| 18263X86 | O Rh+ | 4 | 0 | 0 | 4 | 4 | 0 | 4 | 0 | 2 | 0 | 2 | 0 | 3 | 3 | 3 | 4 | 0 | 3 | 1 | 0 | 0 |

| | | | | | | | | | | | | | | | | | | | | | | |
|---|---|---|---|---|---|---|---|---|---|---|---|---|---|---|---|---|---|---|---|---|---|---|
| 18263X87 | O Rh+ | 4 | 4 | 0 | 4 | 0 | 0 | 4 | 0 | 2 | 0 | 3 | 0 | 3 | 4 | 0 | 4 | 2 | 0 | 3 | 0 | 0 |
| 18263X88 | A Rh+ | 4 | 4 | 0 | 4 | 2 | 0 | 3 | 0 | 2 | 0 | 3 | 0 | 3 | 4 | 0 | 2 | 2 | 0 | 3 | 0 | 0 |
| 18263X89 | O Rh+ | 4 | 4 | 0 | 4 | 0 | 0 | 4 | 0 | 2 | 0 | 3 | 0 | 3 | 4 | 3 | 4 | 0 | 2 | 2 | 0 | 0 |
| 18263X90 | A Rh+ | 4 | 4 | 4 | 4 | 2 | 0 | 4 | 0 | 2 | 0 | 2 | 0 | 2 | 3 | 3 | 2 | 0 | 2 | 2 | 0 | 0 |
| 18263X91 | O Rh+ | 4 | 4 | 0 | 4 | 3 | 3 | 0 | 0 | 4 | 0 | 3 | 0 | 3 | 4 | 3 | 4 | 0 | 0 | 3 | 0 | 0 |
| 18263X92 | O Rh+ | 4 | 4 | 0 | 4 | 0 | 0 | 3 | 0 | 2 | 0 | 3 | 0 | 3 | 2 | 2 | 3 | 2 | 0 | 2 | 0 | 0 |
| 18263X93 | B Rh+ | 4 | 0 | 0 | 4 | 3 | 0 | 3 | 0 | 2 | 4 | 2 | 0 | 3 | 3 | 3 | 2 | 0 | 0 | 3 | 0 | 0 |
| 18263X94 | O Rh+ | 0 | 4 | 0 | 4 | 3 | 0 | 4 | 0 | 2 | 0 | 3 | 0 | 3 | 4 | 3 | 0 | 4 | 3 | 3 | 0 | 0 |
| 18263X95 | O Rh+ | 0 | 4 | 0 | 4 | 3 | 0 | 4 | 2 | 2 | 0 | 3 | 0 | 3 | 4 | 0 | 0 | 3 | 0 | 3 | 0 | 0 |
| 18263X96 | B Rh+ | 0 | 4 | 0 | 4 | 0 | 0 | 4 | 2 | 2 | 0 | 3 | 0 | 2 | 0 | 3 | 3 | 3 | 0 | 2 | 0 | 0 |
| 18263X97 | B Rh+ | 4 | 4 | 0 | 4 | 0 | 0 | 0 | 0 | 2 | 0 | 2 | 0 | 3 | 4 | 3 | 0 | 3 | 0 | 3 | 0 | 0 |
| 18263X98 | O Rh+ | 0 | 4 | 4 | 4 | 3 | 0 | 4 | 0 | 2 | 0 | 3 | 0 | 3 | 4 | 4 | 3 | 2 | 0 | 2 | 0 | 0 |
| 18263X99 | O Rh+ | 0 | 4 | 0 | 4 | 3 | 0 | 0 | 0 | 2 | 0 | 2 | 0 | 3 | 4 | 3 | 3 | 2 | 2 | 0 | 0 | 0 |
| 18263X100 | O Rh+ | 4 | 0 | 0 | 4 | 4 | 0 | 4 | 0 | 2 | 0 | 2 | 0 | 3 | 2 | 3 | 4 | 0 | 2 | 0 | 0 | 0 |

**Table S2**. Genotypes and serological phenotypes for discordant samples using commonly typed variants to infer blood group phenotypes.

| Sample Count | Blood Group | Phenotype by Serology | Phenotype inferred from genotype | Genotype | Resolved? |
|---|---|---|---|---|---|
| 1 | RHD | D- | D+ | Two Copies of *RHD* | Yes: rs748783394 |
| 1 | RHCE (C/c) | C+c+ | C-c+ | Two Copies of RHCE Exon 2 | No |
| 1 | RHCE (E/E) | E+e+ | E-e+ | rs609320: 0/0 | Yes: rs141398055 |
| 4 | Duffy | Fy(a-b-) | Fy(a-b+) | rs2814778: 0/1 rs12075: 1/1 | Yes |
| 1 | Lewis | Le(a+b+) | Le(a-b+) | rs601338: 0/1 rs1047781: 0/0 rs28362459: 0/0 rs812936: 0/1 rs778986: 0/1 | Candidate: rs373779096 |
| 1 | Lewis | Le(a-b-) | Le(a-b+) | rs601338: 0/0 rs1047781: 0/0 rs28362459: 0/1 rs812936: 0/1 rs778986: 0/1 | Yes: rs3894326 |
| 1 | Lewis | Le(a-b-) | Le(a-b+) | rs601338: 0/1 rs1047781: 0/0 rs28362459: 0/1 rs812936: 0/1 rs778986: 0/1 | Yes: rs3894326 |
| 1 | Lewis | Le(a-b-) | Le(a+b-) | rs601338: 1/1 rs1047781: 0/0 rs28362459: 0/1 rs812936: 0/1 rs778986: 0/1 | Yes: rs3894326 |

| | | | | | |
|---|---|---|---|---|---|
| 1 | Lutheran | In(Lu) | Lu(a-b+) | rs28399653: 0/0 | Candidate: rs533045163 and rs184739796 |
| 5 | MNS (M/N) | M+N+ | M-N+ | rs7682260: 0/0 rs7687256: 0/0 rs7658293: 0/0 | No |
| 1 | MNS (M/N) | M+N+ | M+N- | rs7682260: 1/1 rs7687256: 1/1 rs7658293: 1/1 | No |
| 1 | MNS (S/s) | S-s- | S+s- | rs7683365: 1/1 | Yes: rs139511876, DEL1 |
| 1 | MNS (S/s) | S+s- | S+s+ | rs7683365: 0/1 | Candidate: Dantu SV |
| 1 | MNS (S/s) | S+s+ | S+s- | rs7683365: 1/1 | No |
| 1 | P1 | P2 | P1 | rs66781836: 0/1 rs5751348: 0/1 rs8138197: 0/1 rs2143918: 0/1 | Candidate: *A4GALT* -289A>C |

### References

1. Singleton BK, Green CA, Avent ND, et al. The presence of an RHD pseudogene containing a 37 base pair duplication and a nonsense mutation in africans with the Rh D-negative blood group phenotype. *Blood*. Jan 1 2000;95(1):12-8.
2. Haffener PE, Al-Riyami AZ, Al-Zadjali S, et al. Adaptive admixture at ACKR1 (the Duffy locus) may have shaped Plasmodium vivax prevalence in Oman. *bioRxiv*. 2024:2024.03.06.583766. doi:10.1101/2024.03.06.583766
3. Storry JR, Reid ME, Fetics S, Huang CH. Mutations in GYPB exon 5 drive the S-s-U+(var) phenotype in persons of African descent: implications for transfusion. *Transfusion*. Dec 2003;43(12):1738-47. doi:10.1046/j.0041-1132.2003.00585.x
4. Leffler EM, Band G, Busby GBJ, et al. Resistance to malaria through structural variation of red blood cell invasion receptors. *Science*. Jun 16 2017;356(6343)doi:10.1126/science.aam6393
